# Inhibition of nonsense-mediated decay rescues functional p53β/γ isoforms in MDM2-amplified cancers

**DOI:** 10.1101/2020.04.06.026955

**Authors:** Jayanthi P. Gudikote, Tina Cascone, Alissa Poteete, Piyada Sitthideatphaiboon, Qiuyu Wu, Naoto Morikawa, Fahao Zhang, Shaohua Peng, Pan Tong, Lerong Li, Li Shen, Monique Nilsson, Phillip Jones, Erik P. Sulman, Jing Wang, Jean-Christophe Bourdon, Faye M. Johnson, John V. Heymach

**Affiliations:** Department of Thoracic Head and Neck Medical Oncology, The University of Texas MD Anderson Cancer Center, Houston, Texas; Department of Medicine, Division of Medical Oncology, Chulalongkorn University/King Chulalongkorn Memorial Hospital, Bangkok, Thailand; Department of Melanoma Medical Oncology, The University of Texas MD Anderson Cancer Center, Houston, Texas; Division of Pulmonary Medicine, Allergy, and Rheumatology, Department of Internal Medicine, Iwate Medical University School of Medicine, 19-1, Uchimaru, Morioka Iwate, 020-8505, Japan; Department of Bioinformatics and Computational Biology, The University of Texas MD Anderson Cancer Center, Houston, Texas; Novartis Pharmaceutical Corporation, Fort Worth, Texas; Institute for Applied Cancer Science, The University of Texas MD Anderson Cancer Center, Houston, Texas; Department of Radiation Oncology and Brain and Spine tumor Center, Laura and Isaac Perlmutter Cancer Center, NYU Lagone School of Medicine, New York; The University of Texas MD Anderson Cancer Center Graduate School of Biomedical Sciences, Houston, Texas; School of Medicine, Dundee Cancer Center, Clinical Research Center, University of Dundee, Dundee, Scotland, UK

**Keywords:** HPV-associated cancers, MDM2 amplified cancers, NMD inhibition, p53β and p53γ isoforms, Restoration of p53 functions, Therapeutic strategy for p53 deficient cancers

## Abstract

Common mechanisms for p53 loss in cancer include expression of MDM2 or the human papilloma virus (HPV)-encoded E6 protein which both mediate degradation of wild-type (WT) p53 (p53α). Here, we show that two alternatively-spliced, functional, truncated isoforms of p53 (p53β and p53γ, containing exons 1-9 of the p53 gene) can be markedly upregulated by pharmacologic or genetic inhibition of nonsense mediated decay (NMD), a regulator of aberrant mRNA stability. These isoforms lack the MDM2 binding domain and hence have reduced susceptibility to MDM2-mediated degradation. In MDM2-overexpressing cells bearing wildtype *TP53* gene, NMD blockade increased p53β/γ expression and p53 pathway activation, enhanced radiosensitivity, and inhibited tumor growth. A similar pattern was observed in HPV^+^ cancer cells and in cancer cells with p53 mutations downstream of exon 9. These results identify a novel therapeutic strategy for restoration of p53 function in tumors rendered p53 deficient through MDM2 overexpression, HPV infection, or certain p53 mutations.

## INTRODUCTION

Loss of p53 function, the most common alterations in cancer, occurs via multiple mechanisms including *TP53* gene mutations or degradation of WT p53 proteins mediated by MDM2 and HPV-E6 (Vousden and Lu 2002, Levine 2019). The development of therapeutic approaches for restoration of p53 tumor suppressive function is therefore a critical unmet need. Approaches currently under investigation in WT TP53 tumors, include inhibitors of MDM2 and gene therapy (Merkel, Taylor et al. 2017), although none have yet been validated clinically. *TP53* encodes twelve distinct isoforms (Bourdon, Fernandes et al. 2005) among which, p53β and p53γ retain key functions of wild-type p53 (Bourdon, Fernandes et al. 2005, Marcel, Fernandes et al. 2014). Unlike full-length p53α, these isoforms lack the C-terminal negative regulatory region containing protein degradation signals (ubiquitinated lysines at positions 370, 372, 373, 381, 382 and 386 (Rodriguez, Desterro et al. 2000, Anczukow, Ware et al. 2008, Poyurovsky, Katz et al. 2010, Laptenko, Tong et al. 2016)) and hence, are less susceptible for MDM2-mediated degradation (Camus, Menendez et al. 2012). However, because these isoforms are generated by alternative splicing of intron-9, resulting in premature termination codons (PTCs) in exon 9β or exon 9γ, they are likely to be degraded by nonsense-mediated decay (NMD), a RNA surveillance pathway that degrades transcripts with PTCs-arising from nonsense mutations, errors in transcription/splicing, alternative splicing and gene rearrangements (Lewis, Green et al. 2003, Chang, Imam et al. 2007, Hwang and Maquat 2011). Consistent with this, p53β isoform was shown to be NMD susceptible (Anczukow, Ware et al. 2008, Cowen and Tang 2017). Interestingly, though p53γ has similar NMD triggering features as p53β, its NMD susceptibility has not been shown.

In this report, we explored whether the endogenous p53 isoforms content can be manipulated by altering processes involved in mRNA degradation in MDM2-overexpressing and HPV-associated cancer cells to restore the p53-induced cell death pathway abrogated by the overexpression of MDM2 or HPV-E6. We hypothesized that NMD inhibition can increase the expression of p53β/γ isoforms promoting p53-induced cell death pathway and that this strategy can be used to overcome the abrogation of the p53 induced cell death pathway by MDM2-overexpression, HPV infection, or NMD-inducing p53 mutations downstream of exon 9, alterations which together comprise approximately 8% of p53-deficient tumors (The Cancer Genome Atlas (TCGA), (Arbyn, de Sanjose et al. 2012, Campbell, Alexandrov et al. 2016, Zehir, Benayed et al. 2017, Siegel, Miller et al. 2019)) (Tables EV1, EV3 and EV4).

## RESULTS

### NMD inhibition stabilizes p53β/γ isoforms and activates the p53 pathway in MDM2-overexpressing cancer cells

MDM2 is amplified in about 3.7 percent of all cancers, 4 percent of lung cancers and 8 percent of glioblastoma multiforme (GBM), inhibiting p53 tumor suppressive activity (Table EV1). The p53 isoforms modulate p53 transcriptional activity (Bourdon, Fernandes et al. 2005). p53 isoforms expression is regulated by alternative splicing, a process often regulated by NMD. To investigate whether NMD inhibition stabilizes the mRNA expression of p53β/γ isoforms and generate p53β/γ proteins lacking the negative regulatory region (Fig 1A, B), we used non-small cell lung cancer (NSCLC) (H460, H1944 and A549) and GBM (GSC289 and GSC231) cell lines bearing WT *TP53*. Endogenous MDM2 is overexpressed in H1944 and A549 relative to H460, and in GSC231 relative to GSC289 (Fig EV 1A, B). We treated cells with the NMD inhibitor, IACS14140 (compound 11j (Gopalsamy, Bennett et al. 2012), henceforth referred to as NMDi), which selectively inhibits the phosphorylation of UPF1 by SMG1, a critical step for NMD (Chang, Imam et al. 2007, Gopalsamy, Bennett et al. 2012, Kim and Maquat 2019). NMDi treatment (1μM) and subsequent mRNA expression analysis using isoform-specific primers (Table EV2) indicated a significant increase in p53mRNA variants containing the exon 9β in all cell lines, while p53mRNA variants containing the exon 9γ was upregulated in the majority (Fig 1C, D). Moreover, NMDi treatment induced a truncated p53 protein of approximately 47kda-48kda, consistent with the predicted size of p53β/γ, in addition to the WT p53 (Fig 1E). mRNA decay analysis indicated that NMD inhibition prolonged the decay of both p53β and p53γ but not p53α transcript, further confirming the NMD susceptibility of these isoforms (Fig EV2).

**Fig 1.**
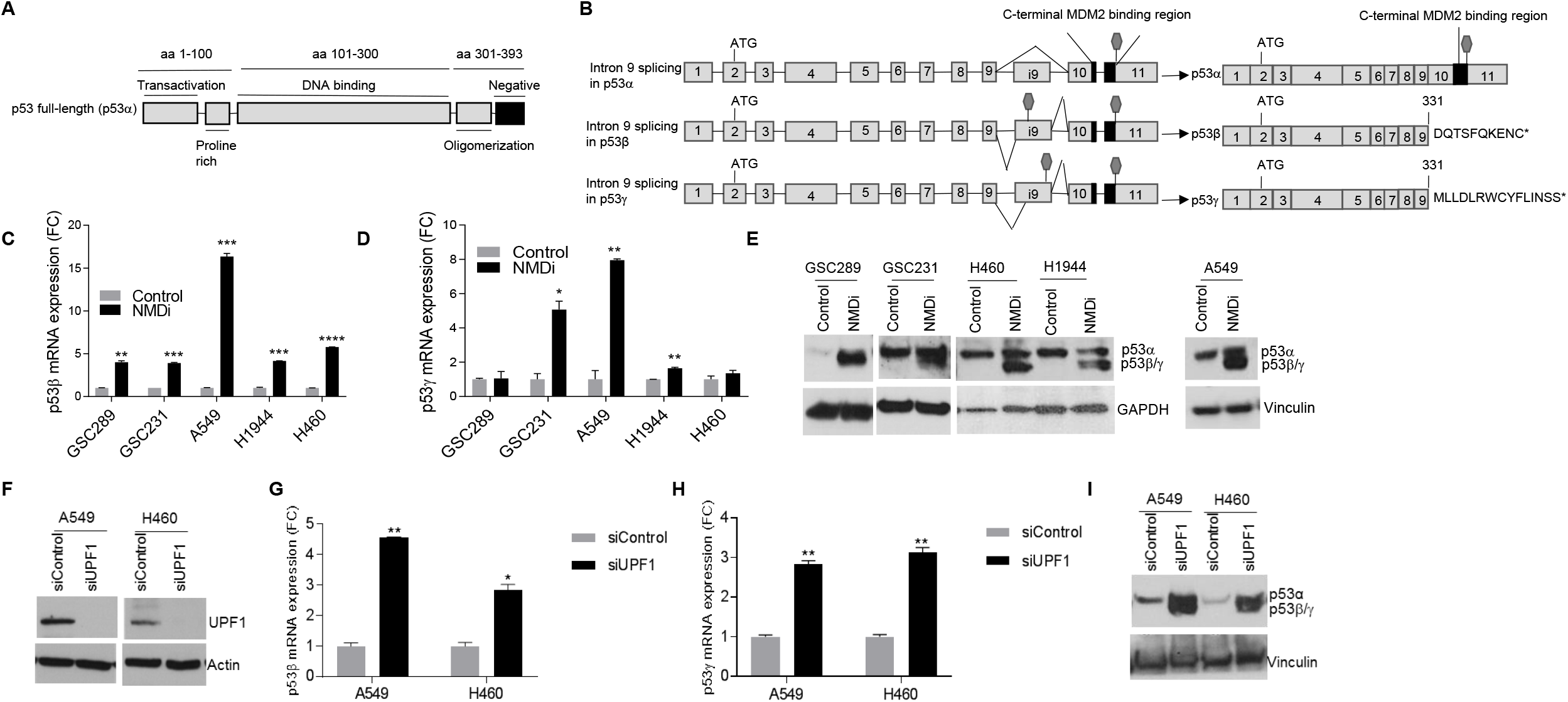
NMD inhibition induces p53β/γ expression in MDM2 overexpressing cancer cells harboring WT *TP53*. (A) Schematic showing functional domains (black lines) in *TP53* gene; aa represents amino acids spanning each domain. (B) Schematic representation of p53α and β /γ isoforms generated by alternative splicing of intron-9. C-terminal MDM2 binding region is depicted in black. Stop signs in exons i9 indicate PTCs acquired by alternative splicing. Amino acids shown at the C-terminus of p53β and p53γ are derived from intron 9. (C and D) Fold change (FC) in the mRNA expression of p53β (C) and p53γ (D) upon NMDi treatment in the shown cell lines. Cells were treated either with DMSO (control) or with 1μM NMDi for 16 hours. RT-qPCR analysis shown (n = 2 technical repeats) are representitives of 3 independent experiments. Mean ± s.e., p values, two tailed t-tests, *≤0.05, **≤ 0.01, ***< 0.001. (E) Western blots showing truncated p53 induction upon NMD inhibition in the cell lines shown. Cells were treated either with DMSO (control) or with 1μM NMDi for 16 hours. GAPDH or Vinculin was used as loading control. One experiment representitive of three is shown. (F) Western blots showing UPF1 knockdown efficiency in A549 and H460 cell lines treated with either sicontrol or siUPF1. Data shown is a representative of two experiments. (G and H) mRNA expression FC of p53β (G) and p53γ (H) respectively in A549 and H460 treated with the indicated siRNAs. RT-qPCR analysis shown (n = 2 technical repeats) are representitives of two independent experiments. Mean ± s.e., p values, two tailed t-tests, *≤0.05, **< 0.01. (I) Western blots showing induction of truncated p53β/γ protein in A549 and H460 cell lines treated with either sicontrol or siUPF1. Data shown is a representative of two experiments.

To gain additional evidence about the NMD susceptibility of p53β and p53γ, we depleted UPF1, a key NMD factor (Chang, Imam et al. 2007, Kim and Maquat 2019), in A549 and H460 cells and analyzed the expression of exon 9β and exon 9γ containing mRNA variant. UPF1 depletion significantly increased the expression of these isoforms (Fig 1F-I). To investigate the effect of NMD inhibition on the p53 pathway, we assessed the mRNA levels of p53 transcriptional targets GADD45A, p21 and PUMA. We observed a significant increase in the mRNA levels of GADD45A and PUMA following NMDi treatment in all five cell lines, while p21mRNA was marginally induced in majority of them, suggesting that inhibition of NMD would activate some p53-inducible promoters but not others (Fig 2A-C). The protein expression level of Puma is strongly induced in A549 and GSC289, while p21 protein is induced only in A549 but not in GSC289 cells (Fig 2D). To test whether NMD inhibition increases the transcriptional activity of p53 on the PUMA promoter, we transfected the luciferase gene reporter construct driven by the PUMA promoter containing p53 binding sites (Yu, Zhang et al. 2001) and assessed the luciferase activity with or without inhibiting NMD in A549 cells. Our results demonstrated a significant increase in the luciferase activity of NMD inhibited cells, indicating that NMD inhibition increased p53 transcriptional activity on PUMA promoter (Fig 2E). UPF1 depleted cells showed increased GADD45A and p21 mRNA expression and increased Puma and p21 protein expression (Fig 2F-H). Taken together, these results suggest that NMD inhibition upregulates alternatively spliced p53 isoforms including the p53β and p53γ, which would change the transcriptional activity of p53 on a subset of the p53-inducible genes in MDM2-overexpressing cancer cells bearing WT *TP53*.

**Fig 2.**
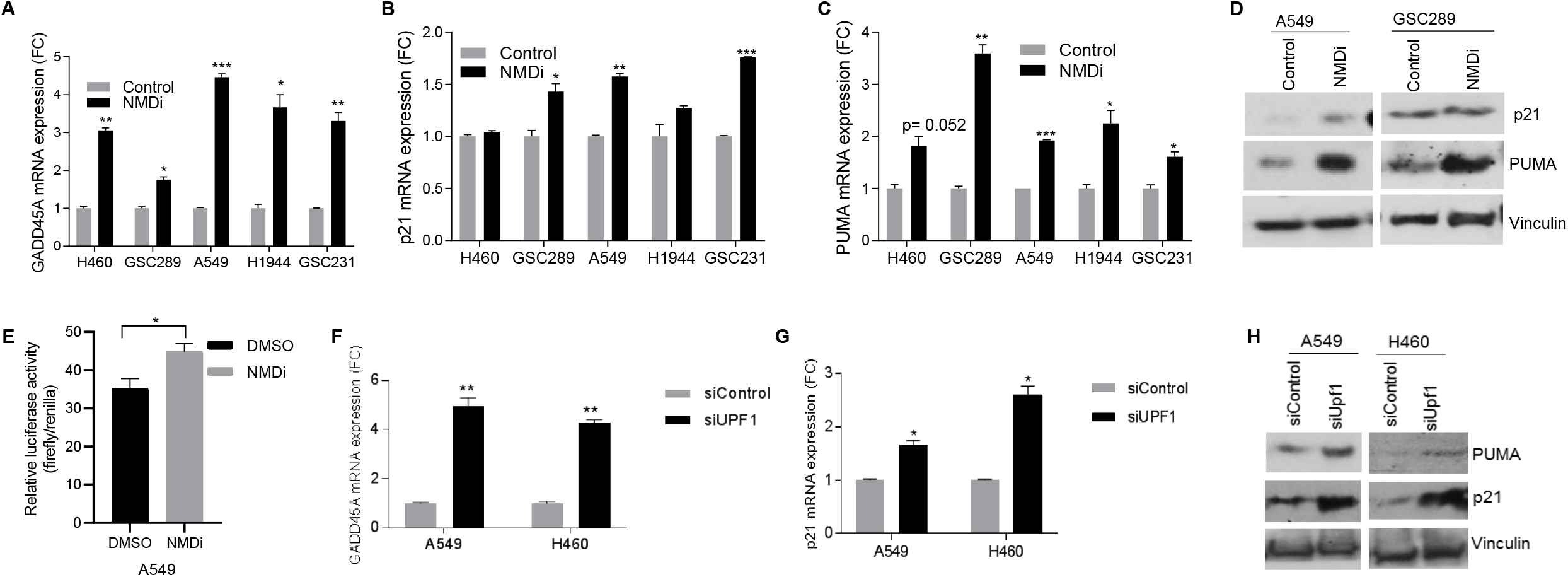
NMD inhibition activates p53 pathway in MDM2 overexpressing cancer cells bearing WT *TP53* gene. (A, B and C) FC in the mRNA expression of GADD45A (A), p21 (B) and PUMA (C) upon NMDi treatment in the shown cell lines. NMDi treatment performed as described in Fig 1. RT-qPCR analysis shown (n = 2 technical repeats) are representitives of 3 independent experiments. Mean ± s.e., p values, two tailed t-tests, *≤0.05, **< 0.01, ***< 0.001. (D) Western blot showing p53 pathway activation in NMDi treated A549 and GSC289. Data shown is a representative of three experiments. (E) NMD inhibition increases p53 transcriptional activity. A549 cells were transfected with 4XBS2-WT luciferase reporter construct and treated with 1.5 μM NMDi for 6 hrs. Luciferase assay was performed as described in the Methods section. Data shown is the mean ± s.e. of three independent transfection experiments. p values, two tailed t-tests, *≤0.05. (F and G) mRNA expression FC of p53 transcriptional target GADD45A (F) and p21 (G) in A549 and H460 treated with the indicated siRNAs. RT-qPCR analysis shown (n = 2 technical repeats) are representitives of two independent experiments. Mean ± s.e., p values, two tailed t-tests, *≤0.05, **< 0.01. (H) Western analysis showing increased p21 and puma expression in A549 and H460 cells treated with the indicated siRNAs. Data shown is representative of two independent experiments.

### Relative contribution of p53γ is higher than that of p53 β in NMD inhibition-induced p53 pathway re-activation

To test whether the effect of NMD inhibition on p21, *PUMA* and *GADD45A* gene expression is dependent on p53 and its isoforms, we compared the expression of p21, PUMA and GADD45A in response to NMD inhibition in A549 cells depleted of all p53 proteins or specific p53 isoform by RNAi-mediated knockdown using validated p53 isoform specific siRNAs. We observed that p21, *GADD45A* and *PUMA* genes were induced differently depending on the co-expressed p53 isoforms, indicating that the observed p53 pathway activation conferred by NMD inhibition is p53-dependent (Fig 3 A-G).

**Fig 3.**
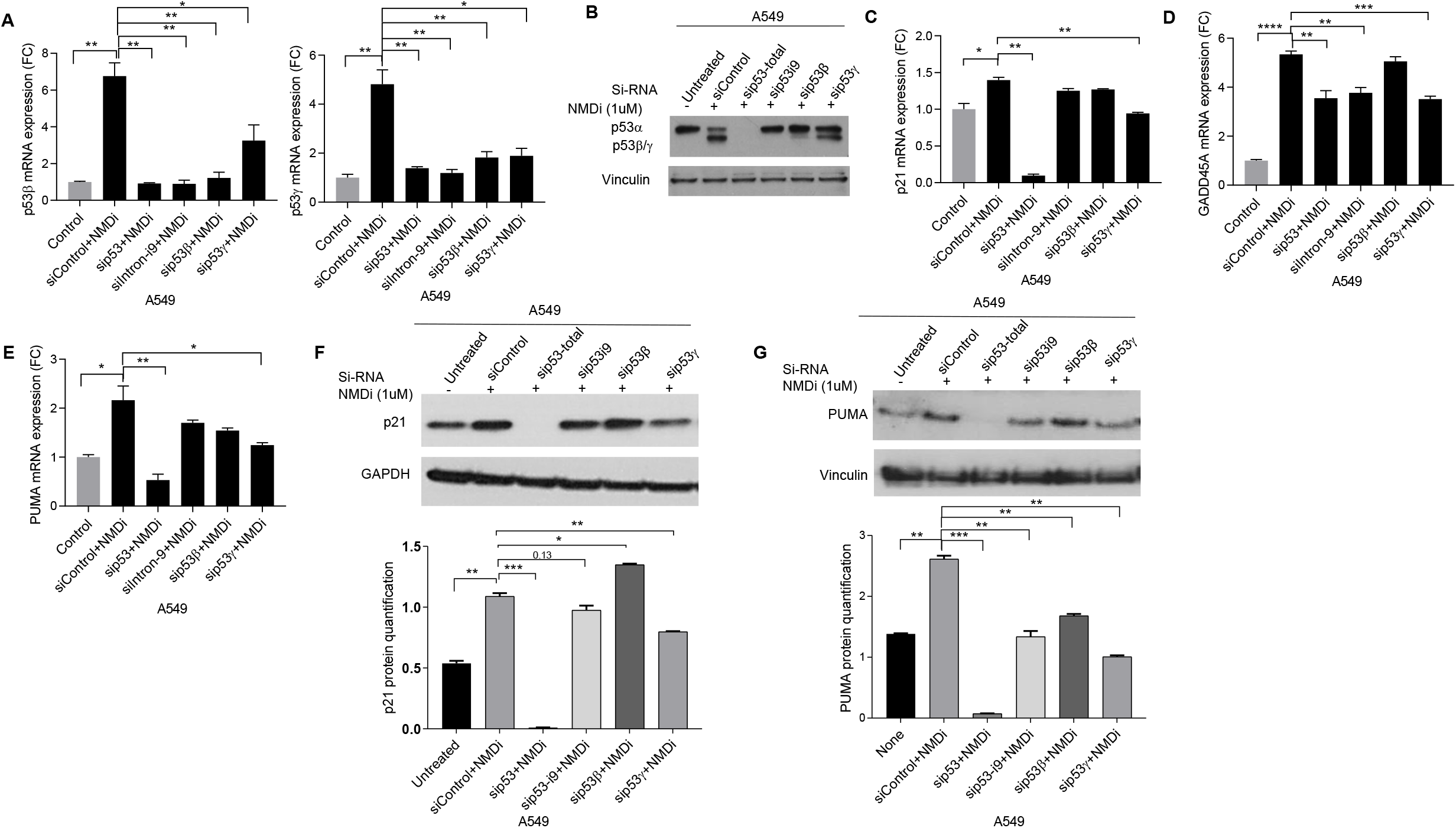
p53 pathway activation conferred by NMD inhibition is p53 dependent and p53γ promotes it. (A) p53β and p53γ mRNA expression FC upon NMD inhibition in A549 cells treated with indicated siRNAs. Cells treated with siRNAs shown were treated with either DMSO (control) or 1μM NMDi, for 16 hours. RT-qPCR (n = 2 technical repeats) shown are representatives of 3 independent experiments. Mean ± s.e., p values, two tailed t-tests, *≤0.05, **< 0.01 (B) Western analysis showing that NMDi-induced truncated p53 proteins are p53β and p53γ isoforms. siRNA and NMDi treatment were done as in (A). Western analysis shown are representatives of 3 independent experiments. (C, D and E) mRNA expression FC of p53 transcriptional targets p21 (C), GADD45A (D) and PUMA (E upon NMD inhibition in A549 cells treated with the indicated siRNAs. siRNA and NMDi treatment were done as in (A). RT-qPCR (n = 2 technical repeats) shown are representatives of 3 independent experiments. Mean ± s.e., p values, two tailed t-tests, *≤0.05, **< 0.01, ***< 0.001, ****< 0.0001. (F and G Western analysis showing p21 (F) and PUMA (G) expression upon NMD inhibition in A549 cells treated with the siRNAs shown. siRNA and NMDi treatment were done as in (A). Lower panels in F and G are the quantifications (2 technical repeats) of the Western blots, normalized to either GAPDH or Vinculin. Western analysis shown are representatives of 3 independent experiments. Mean ± s.e., p values, two tailed t-tests, *≤0.05, **< 0.01, ***< 0.001.

Intron-9 knockdown caused a significant reduction in p53β and p53γ mRNA ((Fig 3A) and truncated p53 protein (Fig 3B) confirming the requirement of intron-9 for NMDi-induced upregulation of p53β and p53γ. siRNAs specific to p53β and p53γ also reduced the NMDi-induced expression of these isoforms, confirming their identity (Fig 3A, B). siRNA-targeting of p53β reduced the expression of p53γ and *vice versa* suggesting that their expression may be codependent (Fig 3A). Intriguingly, among the two isoforms, p53γ knockdown showed the greatest reduction in the expression of p53 transcriptional targets (Fig 3C-E). Western analysis corroborated with the mRNA expression data (Fig 3F, G), providing further evidence that p53β/γ isoforms promote p53 pathway activation and that the relative contribution of p53γ appears to be greater than that of p53β in these tumor cells.

### NMD inhibition stabilizes p53β/γ isoforms and re-activates the p53 pathway in HPV positive cancer cells

An estimated 1.8 percent of all cancers are caused by HPV16/18 infection (Table EV3) (Arbyn, de Sanjose et al. 2012, Siegel, Miller et al. 2019). HPV-associated cancers include cervical, head and neck, vulval, vaginal, penile and anal cancers (Table EV3). The major cause of pathogenesis in cancers induced by HPV infection is the p53 deficiency arising from its E6 oncoviral protein-mediated degradation, even though p53 is predominantly WT (Scheffner, Werness et al. 1990, Balz, Scheckenbach et al. 2003, Strati and Lambert 2008, Sano and Oridate 2016, Faraji, Zaidi et al. 2017). The amino acid residues 94-292 in p53 comprise the core binding region for E6-mediated degradation (Li and Coffino 1996, Martinez-Zapien, Ruiz et al. 2016) and though p53β/γ isoforms retain this region, lack of the C-terminus may alter the tertiary structure of the E6/E6AP/p53β/γ complex, impeding their degradation. Given our findings that NMD inhibition induces p53β and p53γ, and promotes re-activation of the p53 pathway in MDM2-overexpressing cells, we examined whether targeting NMD induces a similar response in HPV^+^ cells bearing WT p53 and hence serve as a potential therapeutic approach for these tumors. We used six HPV^+^ head and neck squamous cell carcinoma (HNSCC) cell lines, two of which (HMS001 and UPCISCC090) also showed *MDM2*-overexpression (Fig EV1C). NMDi significantly increased mRNA levels of both β and γ mRNA variant and induced the expression of the corresponding p53β and p53γ proteins (Fig 4A-C). RNAi-mediated knockdown in HMS001 further confirmed the identity of p53β and p53γ (Fig EV3A, B). NMDi significantly increased the expression of p53 transcriptional targets in these cells (Fig 4D-F). Western blot analysis showed increased expression of p21 and puma proteins in NMD-inhibited cells (Fig 4G, H). Evaluation of p53 binding to *PUMA* promoter indicated a significant increase in the p53 binding activity upon NMD inhibition (Fig 4I). Isoform-specific knockdown in HMS001 indicated that p53γ contributes to the increased expression of p21 and PUMA upon NMD inhibition (FigEV3C-D). These results indicate that NMD inhibition effectively reactivates the p53 pathway in HPV^+^ HNSCC regardless of MDM2 status and that this reactivation is dependent at least in part on the p53γ isoform.

**Fig 4.**
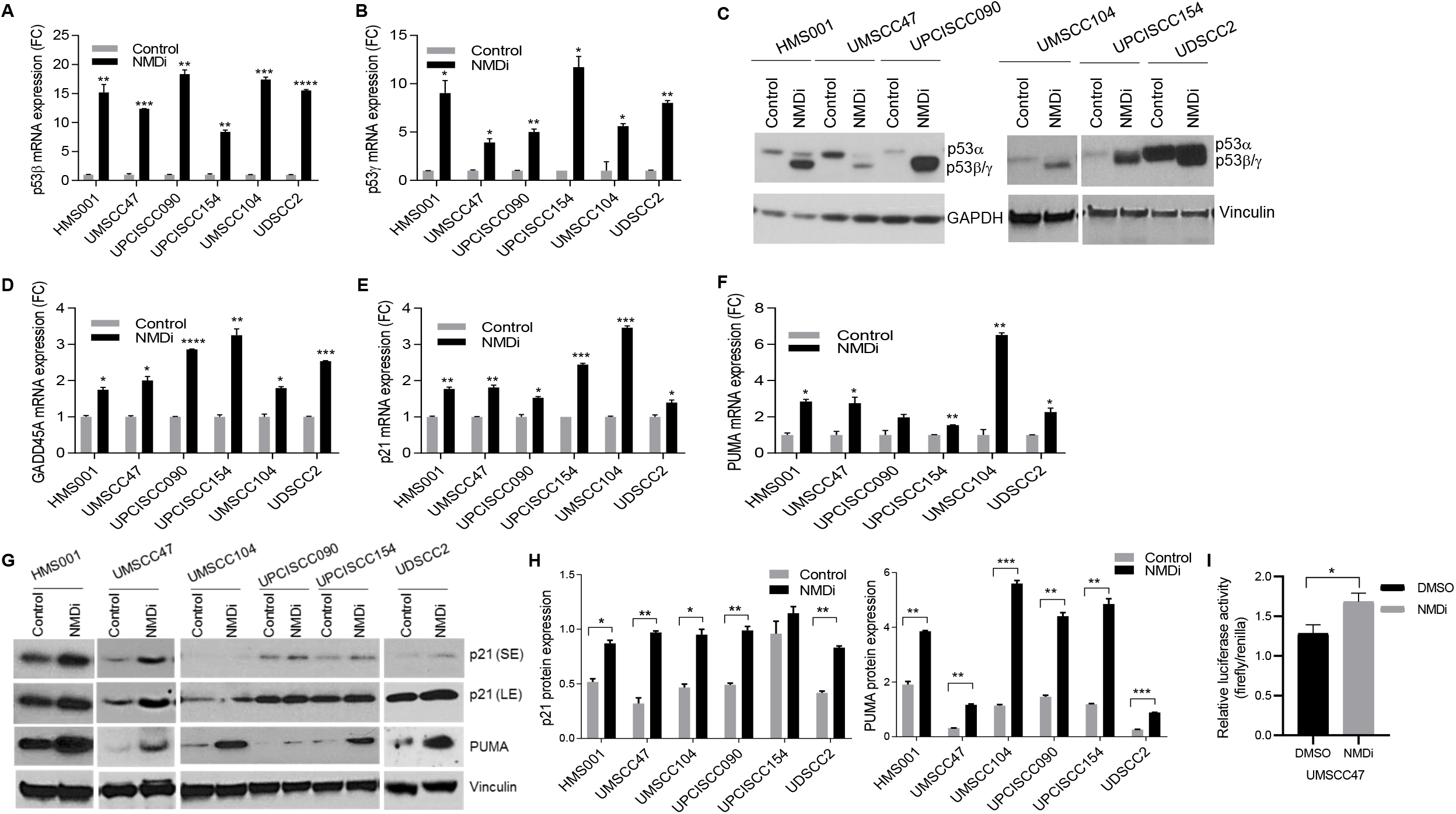
NMD inhibition induces p53β/γ expression and activates p53 pathway in HPV^+^ HNSCC cancer cells. (A and B) mRNA expression FC of p53β (A) and p53γ (B) in HPV^+^HNSCC cell lines shown. Cells were treated with either DMSO (control) or 1μM NMDi for 16 hours. RT-qPCR analysis shown (n = 2 technical repeats) are representatives of 3 independent experiments. Mean ± s.e., p values, two tailed t-tests, *≤0.05, **< 0.01, ***< 0.001, ****< 0.0001. (C) Western analysis showing truncated p53 induction in HPV^+^HNSCC cell lines shown, treated with either DMSO (control) or 1μM NMDi for 16 hours. Western analysis shown are representatives of 3 independent experiments. (D, E and F) mRNA expression FC of GADD45A (D), p21 (E) and PUMA (F) in HPV^+^HNSCC cell lines shown, treated with either DMSO (control) or 1μM NMDi for 16 hours. RT-qPCR analysis shown (n = 2 technical repeats) are representatives of 3 independent experiments. Mean ± s.e., p values, two tailed t-tests, *≤0.05, **< 0.01, ***< 0.001. (G) Western analysis showing p53 pathway activation in HPV¾NSCC cell lines shown, treated with either DMSO (control) or 1μM NMDi for 16 hours. SE, short exposure, LE, long exposure. (H) Quantification (2 technical repeats) of the western blots shown in (G). GAPDH or Vinculin was used as loading control. Mean ± s.e., p values, two tailed t-tests, *≤0.05, **< 0.01, ***< 0.001. (I) NMD inhibition increases p53 transcriptional activity. UMSCC47 cells were transfected with 4XBS2-WT luciferase reporter construct and treated with 1μM NMDi for 16 hrs. Luciferase assay was performed as described in the Methods section. Data shown is the mean ± s.e. of three independent transfection experiments. p values, two tailed t-tests, *≤0.05.

### NMD inhibition restores the p53 pathway and triggers p53β/γ expression in p53 mutant cancer cells

Somatic mutations in *TP53* are found in about 45 percent of all cancers and of these, 34 percent are truncating mutations (Table EV4) that lead to p53 deficiency. About 4.8 percent of the truncating mutations are located either at the end of exon 9 (codon 331) or downstream of it (Table EV 4). If p53 transcripts bearing PTC-generating mutations at the C-terminal end are protected from degradation by NMD, they could produce near full-length protein and rescue p53 function (Fig EV4A, B). To test this, we utilized H2228 (NSCLC), TCCSUP (urinary bladder cancer) and UACC-893 (breast cancer) cells harboring PTCs in p53 exons 9 (H2228) and 10 (TCCSUP and UACC-893) respectively (Fig 5A). NMDi increased the expression of mutant p53 mRNA and protein (Fig 5B, C), consistent with the earlier studies reporting the rescue of nonsense mutation-bearing p53 transcripts by NMD inhibition (Floquet, Deforges et al. 2011, Zhang, Heldin et al. 2017). In addition, NMDi increased the expression of p53β/γ transcripts (Fig 5D, E) and significantly increased mRNA and protein levels of p53 transcriptional targets (Fig 5F-I). These results indicate that for TP53 mutations resulting in PTCs in C-terminal exons, NMD inhibition may help restore p53 function by two distinct mechanisms: preventing the degradation of the mutant transcript, and enhancing the expression of functional p53β/γ isoforms which are truncated upstream of the PTC-inducing mutation. Moreover, in the case of cancers bearing missense mutations downstream of p53 exon 9 resulting in loss of function (LOF) or gain of function (GOF) (codon 331; Table EV4), NMD inhibition could restore p53 function by upregulating p53β/γ isoforms that lack the canonical C-terminus and hence the mutations (Fig EV4C, D).

**Fig 5.**
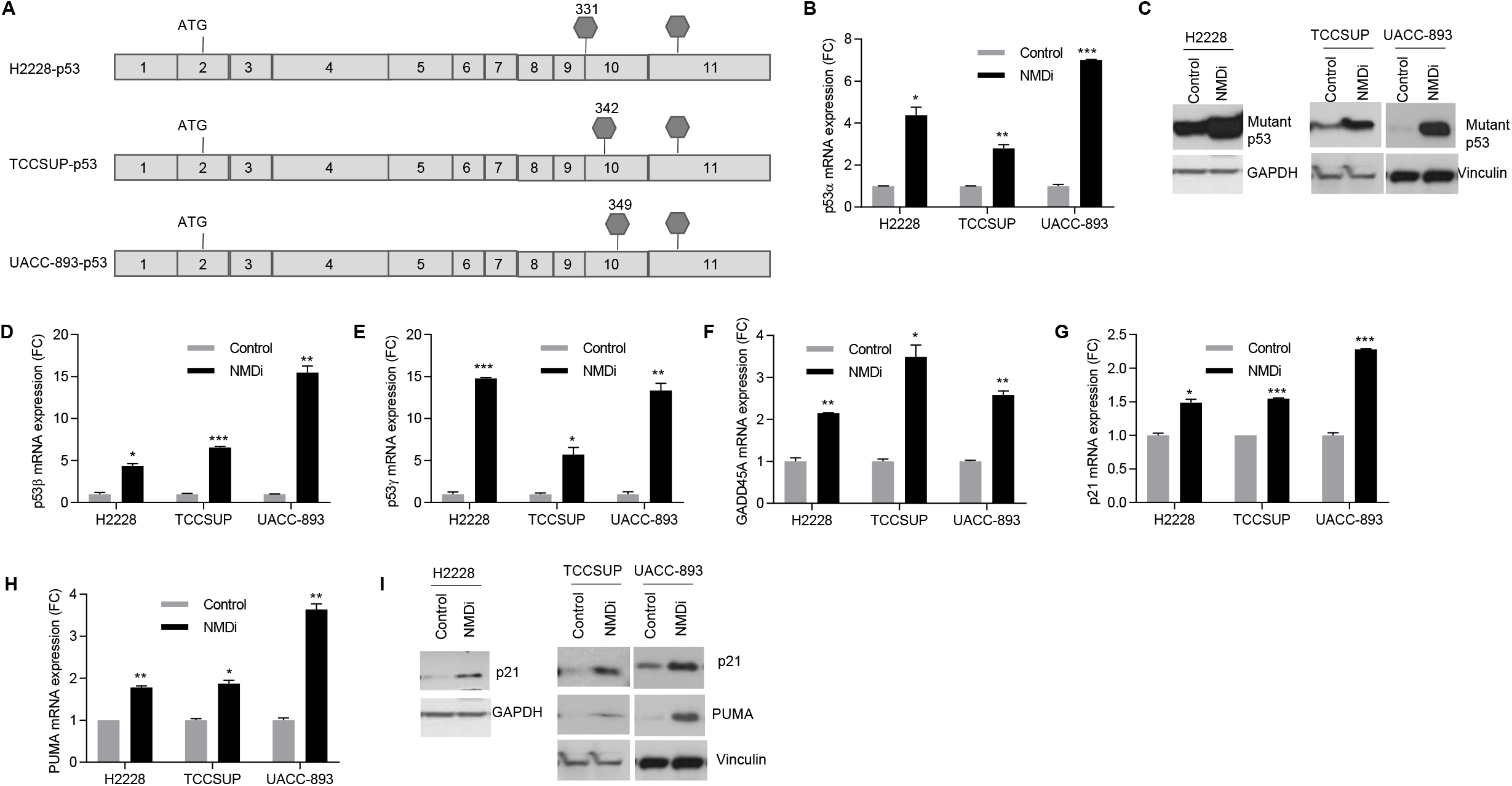
NMD inhibition restores the p53 pathway and triggers p53β/γ expression in p53 mutant cancer cells. (A) Schematic showing the location of PTCs in p53 mutant cell lines. (B) mRNA expression FC of total p53 in p53 mutant cells treated with either DMSO or 1μM NMDi for 16 hours. Data shown (n = 2 technical replicates) are representitives of 3 independent experiments. Mean ±s.e., p values, two tailed t-tests, *≤0.05, **< 0.01, ***< 0.001. (C) Western blots showing increased expression of truncated p53 upon NMDi treatment in p53 mutant cells. Cells were treated as in (A). (D and E) mRNA expression FC of p53β (D) and p53γ (E) isoforms in NMDi treated p53 mutant cells. Cells were treated as in (A). Data shown (n = 2 technical replicates) are representitives of 3 independent experiments. Mean ±s.e., p values, two tailed t-tests, *≤0.05, **< 0.01, ***< 0.001. (F, G and H) mRNA expression FC of p53 transcriptional targets GADD45A (F), p21 (G) and PUMA (h). Cells were treated as in (A). Data shown (n = 2 technical replicates) are representitives of 3 independent experiments. Mean ±s.e., p values, two tailed t-tests, *≤0.05, **< 0.01, ***< 0.001. (I) Western blots showing increased p21 and PUMA protein expression upon NMD inhibition. Cells were treated as in (A). GAPDH or vinculin were used as loading controls for Western blots.

### NMD inhibition induces apoptosis, reduces tumor cell viability and enhances radiosensitivity

p53 activation confers radiosensitivity and chemosensitivity in cancer cells (Fei and El-Deiry 2003, Lu and El-Deiry 2009). Our finding that NMD inhibition re-activated the p53 pathway prompted us to evaluate whether NSCLC, HPV^+^ HNSCC and GBM cell lines are sensitive to NMDi and whether NMDi enhances radiation sensitivity. Our results indicated that NSCLC, HPV^+^ HNSCC and GBM cell lines are sensitive to NMDi (Fig EV5A). Moreover, majority of the HPV^+^ HNSCC cell lines also showed higher sensitivity to NMDi compared to nutlin, a widely used small molecule MDM2-specific inhibitor (Merkel, Taylor et al. 2017), and chemotherapy agents, cisplatin, etoposide and pemetrexed (Fig EV5B).

Because Puma and p21 involved in apoptosis and cell-cycle arrest respectively, were upregulated by NMD inhibition, we also performed apoptosis assay and cell cycle analysis. Analysis of NMDi treated cells for apoptosis by Annexin V FITC/7AAD staining indicated a significant increase in both early and late apoptotic cell population as compared to their respective controls (Fig 6A-E). Cell cycle analysis showed increased G0/G1 and decreased G2/M populations in NMDi treated cells, indicating that NMD inhibition disrupts cell cycle progression (Fig EV5C).We next tested the effect of NMD inhibition on colony forming ability. Results showed that NMDi treatment significantly reduced the colony-forming ability of NSCLC and HPV+ HNSCC cells (Fig 7A). UPF1 depletion showed similar results (Fig 7B), further supporting the finding. Since p53 is also implied in increasing the response to ionizing radiation (IR) (Fei and El-Deiry 2003), we next treated NSCLC and HPV^+^HNSCC cells with radiation alone or in combination with NMDi and evaluated cell viability. NMD inhibition increased the radiotherapy sensitivity of NSCLC and HPV^+^HNSCC cells (Fig 7C).

**Fig 6.**
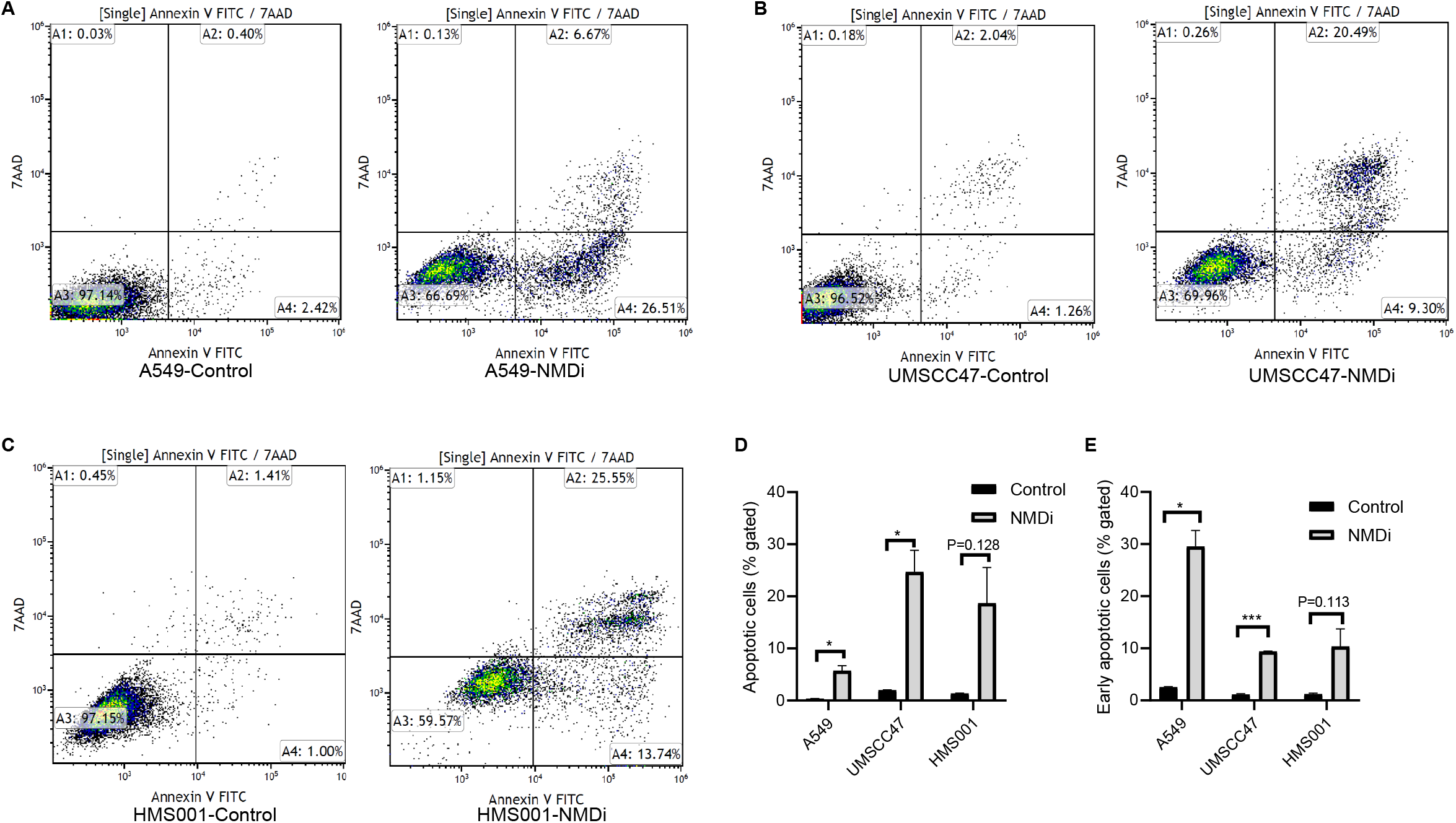
NMD inhibition induces apoptosis in NSCLC and HPV^+^ HNSCC cells. (A, B and C) FACS analysis of either control or NMDi treated A549 (A), UMSCC47 (B) and HMS001 (C) cells. Numbers in quadrant A2 indicate percent of cells that have undergone apoptosis and the numbers in quadrant A4 indicate percent of cells in early apoptotic phase. Data shown are the representitives of two independent experiments. (D and E) Quantification of apoptotic and early apoptotic cells either treated or not treated with NMDi. Data shown are the average of two independent experiments. Mean ±s.e., p values, two tailed t-tests, *≤0.05, **< 0.01, ***< 0.001.

**Fig 7.**
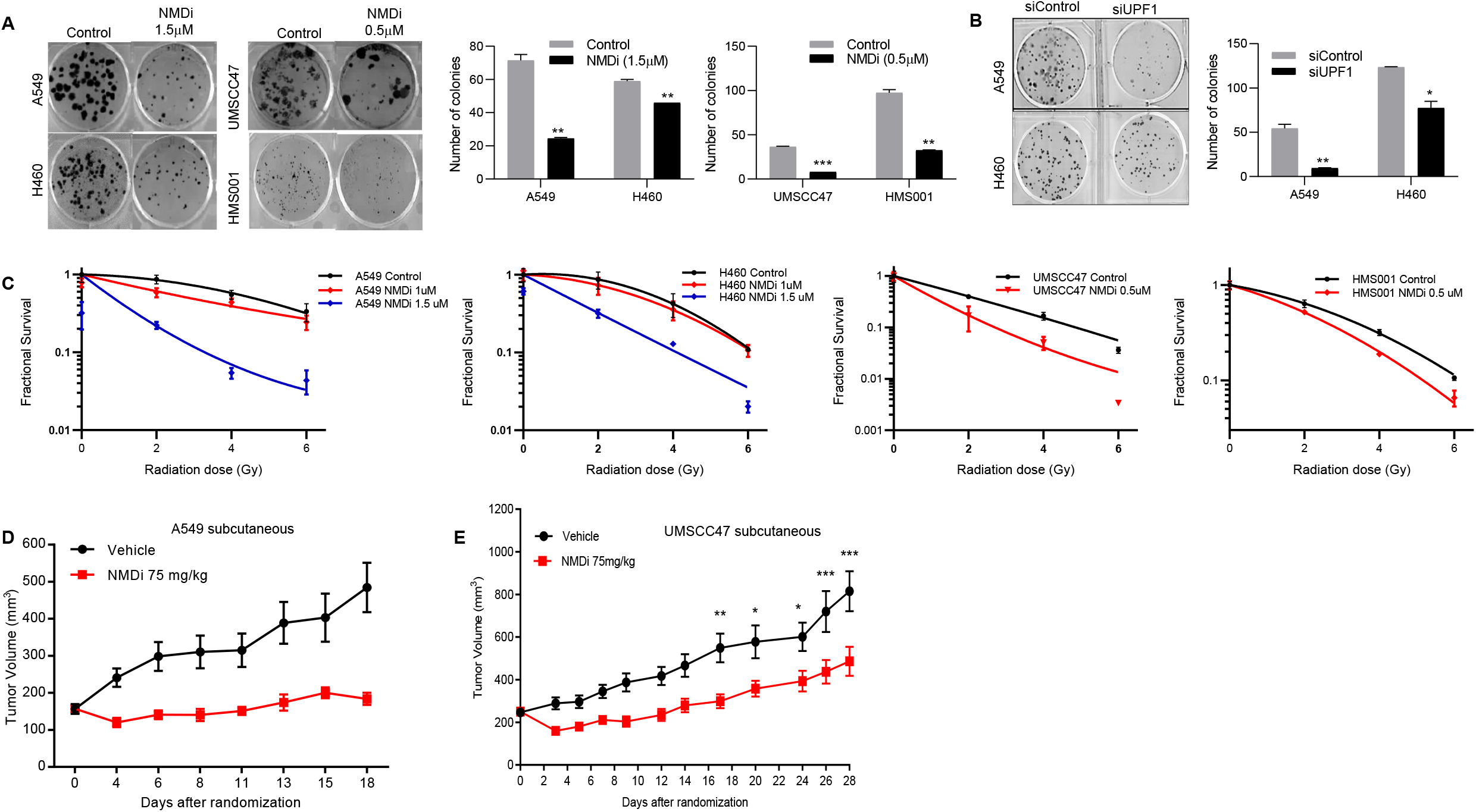
NMD inhibition reduces colony forming ability, increases radiation sensitivity and impairs tumor growth. (A) Clonogenic assay showing reduction in the colony forming ability upon NMD inhibtion (left panels). Quantification (n = 2 experiments) of the colonies from the clonogenic assay (right panels). NSCLC and HPV^+^ HNSCC cells were treated with either DMSO or indicated concentrations of NMDi. Mean ±s.e., p values, two tailed t-tests, *≤0.05, **< 0.01, ***< 0.001. Data shown is the representitive of two experiments. (B) Clonogenic assay (left panel) showing reduction in the colony forming ability upon UPF1 depletion. Cells were treated with either sicontrol or siRNA against UPF1, 24 hours later plated for clonogenic assay. Data shown is the representitive of two experiments. Right panel: quantification (n=2 experiments) of the colonies from respective treatments. Mean ±s.e., p values, two tailed t-tests, *≤0.05, **< 0.01. (C) NMD inhibition increases radiation sensitivity. Clonogenic survival assay of NSCLC and HPV^+^HNSCC cell lines. Cells were treated either with DMSO (control) or with indicated concentrations of NMDi for sixteen hours and exposed to different doses of radiation. Fractional survival was analyzed as described in Methods. (D and E) Subcutaneous xenograft tumor growth in nude mice using A549 (D) and UMSCC47 (E). Mice were injected with either A549 or UMSCC47 cells, randomized as described in Methods. Randomized mice (n=10 per group) were then treated either with vehicle or with 75mg/kg of NMDi for indicated number of days on a five days of treatment and two days of no treatment regimen.

Collectively, our results indicate that NMD inhibition activates p53 pathway, induces apoptosis, disrupts cell cycle progression leading to reduction in cell viability and colony forming ability and, sensitizes cells to radiation.

### NMD inhibition impairs the growth of NSCLC and HPV+ xenograft tumors

We next evaluated the therapeutic implications of NMD inhibition, by assessing the in vivo growth of A549 and HPV^+^ UMSCC47 tumors. Nude mice were injected subcutaneously with A549 or UMSCC47 cells and randomized to receive vehicle or NMDi. In both tumor models, we observed a significant reduction in tumor volume in NMDi-treated animals, as compared to the vehicle-treated animals (Fig 7D, E). We next tested the effect of NMDi on tumor growth in an orthotopic model of UMSCC47 in which tumor cells were injected into the tongue of nude mice. NMDi treatment significantly inhibited tumor growth (Fig EV5D). To investigate whether NMDi treatment induced the expression of p53β/γ isoforms and activated p53 pathway in vivo, we collected tumor tissues from animals and evaluated expression of p53 isoforms and transcriptional targets by RT-PCR. We observed increased p53β, p53γ, PUMA, GADD45A and BAX mRNA expression in NMDi treated A549 tumors and an increased expression of p53γ, p21, GADD45A and PUMA mRNAs in NMDi treated UMSCC47 tumors, compared to their respective vehicle treated controls (Fig EV5E). These findings imply that NMD inhibition reactivates the p53 pathway in vivo and impairs tumor growth.

## DISCUSSION

p53 is the most frequently inactivated tumor suppressor in cancer. Here, we investigated NMD inhibition as a strategy to re-activate p53 and, in turn, impair tumor growth and sensitize tumor cells to commonly used therapeutic regimens. We provide evidence that NMD inhibition stabilizes p53β and γ isoforms and restores p53 activity in several major types of p53-deficient tumor cells including those with nonsense mutations downstream of exon 9 or MDM2 overexpression as well as HPV^+^ HNSCC cells (Fig 8A-D). p53β and γ isoforms lack the C-terminal negative regulatory region and hence less susceptible for MDM2 and HPV-E6-mediated degradation than p53α. Enhancing p53β and γ isoforms expression may therefore particularly benefit cancers that are caused by degradation of wild-type p53 by its negative regulators (Fig. 8C).

**Fig 8.**
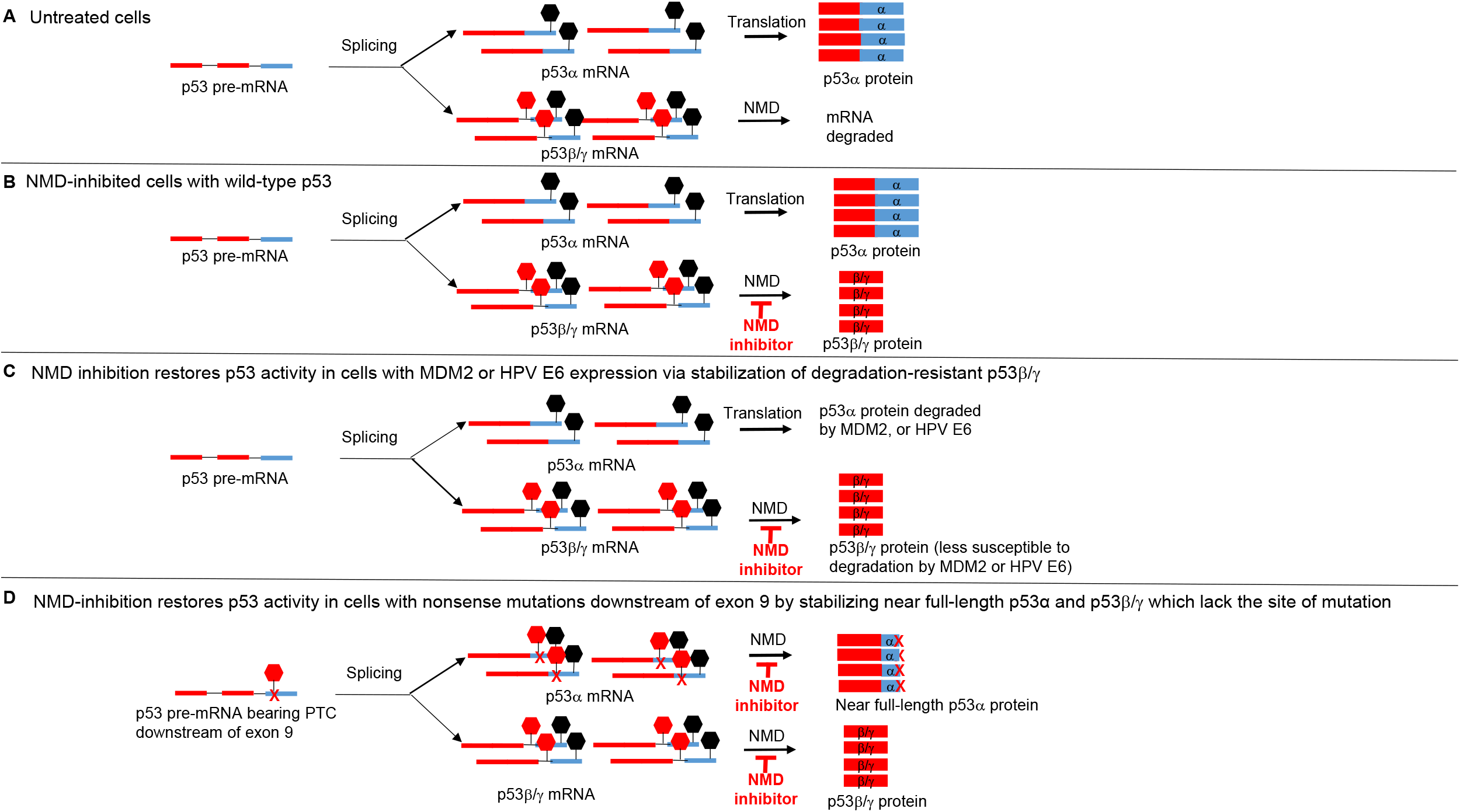
NMD inhibition stabilizes p53β/γ isoforms and restores p53 activity in p53 deficient tumors. (A) In normal cells, p53 pre-mRNA splicing results in p53α and p53β/γ mRNAs. p53β/γ mRNA is degraded by NMD, p53α mRNA is translated. (B) NMD inhibition stabilizes p53β/γ mRNAs which in turn are translated into p53β/γ proteins. (C) In MDM2 amplified or HPV-associated cancers, p53α protein is degraded. NMD inhibition overcomes p53 deficiency by increasing the expression of functional p53β/γ isoforms that are degradation resistant due to the lack of C-terminal negative regulatory region. (D) In p53 mutants bearing nonsense mutations downstream of exon 9. NMD inhibition stabilizes mutant p53 transcripts as well as p53β/γ transcripts, which are translated in to near full-length p53α and p53β/γ proteins respectively. Stop sign in black, normal stop codon; stop sign in red, PTC; x, nonsense mutation downstream of exon 9. p53 exons 1-9 are depicted in red, exons 10 and 11 are depicted in light blue.

NMD inhibition using aminoglycosides or ataluren (PTC124) has been a therapeutic strategy in many diseases caused by nonsense mutations including Duchenne muscular dystrophy and cystic fibrosis (Wilschanski, Yahav et al. 2003, Finkel 2010). These pharmacological NMD inhibitors act by suppressing PTCs caused by nonsense mutations, but are not effective at protecting the degradation of physiological NMD substrates such as alternatively spliced transcripts. The NMDi used in the current study, by contrast, not only efficiently reverses the NMD of the p53 mutant mRNA transcripts but also physiological NMD substrates such as p53β and p53γ, generated by alternative splicing. NMDi treatment enabled the synthesis of near full-length, functional p53 proteins in NSCLC, breast cancer and urinary bladder cancer cell lines bearing nonsense mutations downstream of p53 exon 9. In addition, NMDi also protected p53β and p53γ from degradation in a wide variety of p53-deficient cancer cells including p53 mutant, MDM2 overexpressing and HPV^+^ cell lines. One important NMD triggering feature is the location of a PTC with respect to the final exon-exon junction of an mRNA. In humans, PTCs located at least 50-55 nt upstream of the final exon-exon junction are known to trigger NMD (Kurosaki and Maquat 2016). Even though both p53β and p53γ fulfil this requirement, earlier studies were not able to detect p53γ induction upon NMD inhibition and showed only p53β as an NMD substrate (Anczukow, Ware et al. 2008, Cowen and Tang 2017). In this study, we show that both p53β and p53γ are NMD substrates and NMD inhibition upregulates their expression. Failure to detect p53γ upon NMD inhibition in earlier studies may be because of its low abundance in certain cell types.

p53β and γ isoforms promote transcription of p53 target genes and retain tumor suppressive functions. For instance, p53β and p53γ promote apoptosis and senescence by inducing p53α-dependent transactivation of Bax and p21 respectively (Bourdon, Fernandes et al. 2005, Surget, Khoury et al. 2013, Solomon, Sharon et al. 2014). Moreover, in breast cancer patients, p53β and p53γ levels are associated with more favorable clinical outcome (Bourdon, Khoury et al. 2011, Avery-Kiejda, Morten et al. 2014). p53β expression showed a negative correlation with tumor size and positive correlation with survival in breast cancer patients. High levels of p53β were particularly protective in patients with mutant p53 and may play a role in counteracting the damage inflicted by mutant p53 (Avery-Kiejda, Morten et al. 2014). Furthermore, breast cancer patients expressing p53γ along with mutant p53 had as good a prognosis as those with WT-p53, whereas those who were devoid of p53γ and expressed only mutant p53 showed markedly poor prognosis (Bourdon, Khoury et al. 2011). Our finding that both p53β and γ isoforms can be upregulated by NMD inhibition in p53 mutant cells signifies the therapeutic benefits of NMD inhibition in p53 mutant cancers.

We found that NMD inhibition sensitized NSCLC and HPV^+^HNSCC cells to ionizing radiation (IR). Interestingly, IR has been shown to trigger the expression of p53β, which in turn contributes to IR-induced cellular senescence (Chen, Crutchley et al. 2017). Here, we show that p53β/γ levels are increased by NMD inhibition and that cancer cells in which NMD was inhibited were more sensitive to IR compared to the ones without NMD inhibition. Increased radiation sensitivity in NMD inhibited cells may be because of the protection of both NMD inhibition-induced as well as IR-induced p53β from degradation. Whether p53γ plays a role in enhancing radiation sensitivity needs further investigation.

Drugs developed to target a particular p53 mutation have limited usage against other p53 mutations. For example, NSC31397 restores p53 activity in R175H-p53 mutant cancer cells and p53R3 restores DNA binding of R175H and R273H p53 mutants (Hong, van den Heuvel et al. 2014). The approach described here for p53 restoration could potentially be broadly applied to not only tumors with p53 mutations downstream of exon 9 but also to those that overexpress negative regulators of p53 such as MDM2 and HPV-E6 protein. This novel approach could also be used to eliminate loss of function (LOF) or gain of function (GOF) mutations in about 2.7 percent of all cancers bearing mutations downstream of p53 exon 9 (Table EV4), as p53β/γ isoforms lack the canonical C-terminus (Fig EV4). Collectively, NMD inhibition strategy could benefit approximately 8% of all cancers which bear the appropriate p53 mutations, MDM2 amplification, or HPV E6 expression (MDM2 amplified, 3.7%,; HV16/18-associated cancers, 1.8% and mutations downstream of exon 9, 2.7%: Tables EV1, 3 and 4). The finding that NMD inhibition effectively activates the p53 pathway, increases radiosensitivity, and reduces tumor growth in preclinical models suggests that NMD inhibition merits further investigation as a therapeutic strategy in a wide variety of cancers that are p53-deficient.

## MATERIALS AND METHODS

### Cell lines

A549, H460, H1944, H1395, H2228, TCCSUP and UACC-893 cells were from American Type Culture Collection (ATCC). UMSCC47 and UMSCC104 were from Dr. Thomas E. Carey, (University of Michigan, Michigan) (Brenner, Graham et al. 2010, Tang, Hauff et al. 2012). UDSCC2 was a gift from J. Silvio Gutkind (University of california, San Diego). UPCISCC090 and UPCISCC154 were established at the University of Pittsburgh (White, Weissfeld et al. 2007, Martin, Reshmi et al. 2008). HMS001 cell line was established by Dr. M.L. Gillison’s laboratory (Akagi, Li et al. 2014). GSC231 and GSC289 were generated and finger printed at the laboratories of Dr. Erik Sulman and Dr. Frederick Lang Jr (Department of Neurosurgery, UT-M.D. Anderson Cancer Center). A549, H460 and H2228 were grown in RPMI with 10% FBS. UDSCC2 and UMSCC104 were grown in DMEM containing 10% FBS and 1% nonessential amino acids (NEAA). UPCISCC090 and UPCISCC154 were grown in MEM with 10% FBS and 1% NEAA. UMSCC47 was grown in DMEM with 10% FBS, HMS001 in DMEM/F12 with 10% FBS. HTB5 grown in MEM with 10% FBS, 1% NEAA and 1mM sodium pyruvate. CRL-1902 cells were grown in Lebovitz’s L-15 Medium (ATCC) with 10% FBS. GSC231 and GSC289 cells were grown in Neutral Basal Medium (DMEM/F12 50/50, 1X B27 supplement, 20ng/ml EGF, 20ng/ml bFGF, 1% Pen/Strep solution). All cell lines were negative for mycoplasma.

### NMDi treatment

NMDi was supplied by Dr. Phillip Jones (Institute for Applied Cancer Science, UT-M.D. Anderson Cancer Center, Houston, TX). Cells were treated either with DMSO or with 1μM NMDi for ~16h and harvested for preparing protein lysates and RNA.

### RNAi-mediated knockdown

siRNAs against human UPF1 (aagatgcagttccgctccatttt) (Mendell, ap Rhys et al. 2002), p53β (ggaccagaccagctttc), p53γ (cccttcagatgctacttga) and p53 intron-9 (gaugcuacuugacuuacga) (Marcel, Fernandes et al. 2014) and total p53 (gagguuggcucugacugua, siRNA ID: SASI_Hs02_00302766) were purchased from SIGMA-ALDRICH. 700,000 to 1 million cells were plated in 100mm tissue culture plates overnight. siRNA stocks (100μM) were diluted in RNase-free water, mixed with DharmaFECT 1 (Dharmacon), incubated for 20 minutes at room temperature and added to the cells in medium lacking penicillin/ streptomycin, at a final concentration of 100nM siRNA in a total volume of 8ml/plate. Twenty four hours post transfection, cells were split into two 60mm plates each, for protein and RNA extraction. siUPF1 treated cells were harvested 72 hours post transfection. For sip53, sip53 intron-9, sip53β and sip53γ treated cells, 56 hours post transfection, NMDi was added to a final concentration of 1μM and cells harvested 16 hours post NMDi treatment for protein and RNA extraction.

### Quantitative PCR

RNA was extracted either by using TRI Reagent Soln (Ambion) or RNeasy kit (Qiagen), according to manufacturer’s instructions. Total RNA was quantified using NANO DROP 2000C spectrophotometer (ThermoScientific). cDNA synthesized by reverse transcription using iScript™Reverse Transcription Supermix for RT-qPCR (BIO-RAD), using 1μg of total RNA, according to manufacturer’s instructions. Quantitative PCR (qPCR) was done on cDNA using PerfeCta SYBR Green FastMix Low Rox probes (Quantabio) and the primers listed in Extended Data Table S1. Real-Time PCR done in 7500 Fast Real-Time PCR System (Applied Biosystems). ΔΔ*C*t method was used to calculate fold changes in expression. Human GAPDH was used as an endogenous control for mRNA expression. Fold changes expressed are normalized either to the siControl or DMSO treated controls.

### Assessment of mRNA decay

To assess mRNA decay of p53 isoforms, A549 cells were treated with either DMSO or NMDi (1μM) for 16 hours, transcription was then inhibited by adding 5,6-Dichloro-1-β-D-ribofuranosylbenzimidazole (100μM) and RNA was extracted at various time points. mRNA expression analysis was performed by RT-qPCR.

### Western blot

Protein lysates (30μg-40μg) were resolved on 4%-15% gradient SDS-PAGE gels, transferred to polyvinylidene difluoride (PVDF) membrane using a wet system (BIO-RAD). Membranes were washed briefly in 1X TBS-Tween (TBST) and blocked in 5% (W/V) dried milk in TBST for 1 hour. They were incubated overnight at 4°C in primary antibodies diluted 1:1000 in 5% (W/V) BSA-containing TBST, washed 3 times in TBST and incubated in appropriate horseradish peroxidase-conjugated secondary antibody. Membranes were developed using Super Signal^™^ West Pico PLUS Chemiluminescent Substrate (Thermo Scientific) and exposed to Blue Lite Autorad Film (GeneMate). GAPDH, actin or vinculin was used as loading control. Images shown are representatives of three separate protein isolations and blots run in triplicate. Antibodies used were: p53 (7F5, Catalogue # 2527S), puma (D30C10, Catalogue #12450S), p21 Waf1/Cip1 (DCS60, Catalogue # 2946S), Upf1 (Catalogue # 9435) and GAPDH (Catalogue # 5174) from Cell Signaling Technology, MDM2 (HDM2-323, Catalogue # sc-56154) from Santa Cruz Biotechnology, β-actin (Catalogue # A5441) and Vinculin (Catalogue #V9131) from SIGMA. Quantifications of the protein bands were done using ImageJ (Schneider, Rasband et al. 2012). Protein expressions were normalized to that of either GAPDH, actin or vinculin.

### Luciferase reporter assay

A549 and UMSCC47 cells were plated in 24-well plates and transfected with 1μg/well of 4XBS2 WT reporter construct (Yu, Zhang et al. 2001) (Catalogue # 16593, addgene) using Lipofectamine 2000 (Invitrogen). Renilla construct (25 ng/well, Promega) was co-transfected as a control for transfection efficiency. Approximately 24 hours post transfection, A549 cells were treated with 1.5μM NMDi for 6 hours and approximately 31 hours post transfection, UMSCC47 cells were treated with 1μM NMDi for 16 hours. Cells were harvested in passive lysis buffer and luciferase assay was performed using Dual-Luciferase Assay System (Catalogue # E1960, Promega), according to manufacturer’s instructions. Luminescence was measured using GLOMAX 20/20 Luminometer (Promega). Luciferase activity was normalized with Renilla activity.

### RNAseq, and mutation analysis

RNAseq, HPV status and mutation data were analyzed as described (Kalu, Mazumdar et al. 2017). RNA expression for MDM2 and TP53 across all available cell lines in M.D. Anderson Cancer Center’s Head & Neck (HN) and Lung database were integrated. HPV status was annotated for each HN cell line. The heat map was sorted by RNA expression of MDM2.

PanCancer tumor analysis for MDM2 amplification and TP53 mutation frequency were performed using the data from The Cancer Genome Atlas (TCGA) (Campbell, Alexandrov et al. 2016, Zehir, Benayed et al. 2017) portal.

### Drug treatment for assessing cell viability and IC50

Nutlin 3 (Catalogue# S1061) was purchased from Selleck Chemicals LLC, Houston, TX. Cisplatin, etoposide and pemetrexed were from Institutional Pharmacy, The University of Texas-M.D. Anderson Cancer Center. Six hundred cells/well for H460, 700 cells/well for A549 and 1000 cells/well for all other cell lines were plated in 384-well plates (GreinerBio-one) in triplicates on the same plate. Cells were treated with DMSO or seven different concentrations of serially threefold-diluted drugs, highest concentration being 9.6μM, in a final volume of 40μl. Seventy two hours later, 11 μl of CellTiter Glo (CellTiter-Glo Luminescent Cell Viability Assay, REF G7573, Promega) added, plates shaken for 10 min, luminescence was measured using a FLUOstar OPTIMA microplate reader (BMG LABTECH). Luminescence values were normalized to DMSO-treated cells. Each experiment was repeated two separate times to give biological replicates. IC50 values were estimated using drexplorer software, which fitted multiple dose-response models and identified the best model using residual standard error (Tong, Coombes et al. 2015). In brief, the dose-response data was normalized by the mean response of controls prior to drug parameter estimation. Outlier data points were detected and removed as described in (Tong, Coombes et al. 2015). The best dose-response model was selected based on residual standard error. Drug parameters including inhibitory concentrations IC (IC10~IC90, original dose scale) and area under curve (AUCs, scaled by the curve at response=1; 0 means extremely sensitive and 1 means extremely resistant) were estimated using the drexplorer package (Tong, Coombes et al. 2015). The estimation of IC values were truncated by the observed doses. For example, if IC50 was not achieved, IC50 was estimated as the largest dose used in the experiment. On the contrary, if the drug is highly effective where the lowest dose killed more the 50% of cells, IC50 was represented by the smallest observed dose. Most profiling had two replicates. For these experiments, we assessed reproducibility of the experiments based on concordance correlation coefficient (CCC) and location shift. Experiments with CCC>0.8 or location shift<0.9 were flagged as passing the quality control. A paired t-test was used to compare relative viability between drug treated and DMSO treated control groups.

### Cell cycle analysis

One million cells were plated in 100mm plates. Twenty four hours later, cells were treated either with DMSO or with NMDi (1μM). Cells were harvested 24h and 48h post treatment, fixed in 70% ethanol for 24h at 4°C, washed with 1X phosphate-buffered saline (PBS), treated with 200μg/ml RNase A in 1X PBS for 1 hour at 37°, stained with 40μg/ml propidium iodide (PI) for 20 minutes at room temperature, spun at 500g for 5min to remove PI and the cell pellet was re-suspended in 1XPBS (500μl). Fluorescence-Activated Cell Sorting (FACS) analysis was performed in BDFACSCanto flow cytometer.

### Apoptosis assay

150,000 cells/well for A549 and 300,000 cells/well for UMSCC47 and HMS001 were plated in six well plates. Approximately 24 hours later, cells were treated either with DMSO or with 3μM (A549) or 2 μM (UMSCC47 and HMS001) NMDi for approximately 72 hours. Cells were then trypsinized, washed two times with ice cold PBS and processed for staining using FITC Annexin V Apoptosis Detection Kit with 7AAD (Cat # 640922, BioLegend), according to manufacturer’s instructions. Cells were analyzed by Flow Cytometry.

### Clonogenic survival assay

Exponentially growing cells were plated in duplicates at three dilutions into 6-well plates containing 2 ml medium and incubated for 24 h in a humidified CO2 incubator at 37°C. Subsequently NMDi was added into the medium for 16h. After being pretreated with DMSO (for control) and NMDi, cells were subjected to 0, 2, 4, or 6 Gy of gamma irradiation by using a Mark 1-68A cell irradiator with a Cesium-137 source at a dose rate of 3.254 Gy per minute (J.L. Shepherd & Associates, San Fernando, CA). The medium was then replaced with fresh medium allowing cells to continuously grow for colony formation for 10 to 14 days. To assess clonogenic survival following radiation exposure, cells were fixed with glutaraldehyde (6.0% v/v), stained with crystal violet (0.5% w/v). Colonies with more than 50 cells were counted to determine percent survival and the number of colonies obtained from three replicates was averaged for each treatment. These mean values were corrected according to plating efficiency of respective controls to calculate cell survival for each dose level. After correcting for plating efficiency, the survival fraction was calculated as previously reported (Franken, Rodermond et al. 2006) and fitted to a linear-quadratic model by using SigmaPlot 10.0 (San Jose, CA). Clonogenic survival assays after irradiation were repeated two times for each cell line.

### Assessing colony forming ability

For assessing colony forming ability after UPF1 depletion, cells transfected with either siControl or siUPF1 were plated in duplicate at 200 cells per well concentration, in a six-well plate, approximately 26 hours after transfection. For assessing colony forming ability of NMDi treated cells, cells were plated in duplicate at 100 cells per well concentration in a 6 well plate and 24 hours later, treated with either DMSO or with NMDi. Medium was changed 24 hours post treatment. Ten to fifteen days later, colonies were stained with 0.25% Crystal Violet in 100% methanol. Colonies with more than 50 cells were counted and the number of colonies obtained from three replicates was averaged for each treatment.

### Tumor growth assessment

All animal studies were approved by the Institutional Animal Care and Use Committee at the University of Texas MD Anderson Cancer Center in accordance with National Institutes of Health guidelines. Female mice (Strain 69 Athymic Nude (nu/nu) mice) were purchased from Envigo. Mice were randomly assigned to control or treatment groups. No statistical method was applied to predetermine the sample size and the investigators performing preclinical experiments were not blinded. A549 (5×10^6^ cells per mouse) or UMSCC47 (2×10^6^ cells per mouse) were injected subcutaneously into nude female mice. For orthotopic model, 60,000 cells were injected into the tongue of nude female mice. Mice were randomized for treatment (at least 10 per group) when the tumor size reached ~250-300mm^3^ for the flank model and ~5-8mm^3^ for the orhotopic model. A549 xenografts were treated for 3 weeks, UMSCC47 flank model was treated for 4 weeks and orthotopic UMSCC47 model was treated until the tumor size reached ~30mm^3^. NMDi was administered at the indicated doses on a 5 days on and 2 days off regime, for all three models. Tumors were harvested at the end of the treatment regime and processed for RNA collection and RT-qPCR analysis.

### Statistical Analysis

For assessing cell viability, a paired t-test was used to compare relative viability between drug treated and DMSO treated control groups. For mRNA expression fold change analysis, unpaired t-test (two-tailed) were used to compare the relative expression between drug treated and DMSO treated control groups. For analysis of tumor growth data, quantitative data were subjected to two-way analysis of variance (ANOVA).

## ACKNOWLEDGEMENTS

We thank Irene Guijarro Munoz for editorial assistance and Huiying Sun for the technical assistance.

## AUTHORS CONTRIBUTIONS

J.P.G. designed and performed the experiments and analyzed the data. J.P.G. wrote the manuscript with the assistance of all other authors. T.C., A.P., F.Z., Q.W. and S.P performed the *in vivo* mouse experiments. T.C. and A.P. analyzed the tumor growth. P.S. performed radiation sensitivity assays. N.M. helped with drug treatments and analysis. P.T., L.L., L.S. and J.W. performed the statistical analysis. M.N. helped with the manuscript construction and writing. P.J. provided the NMDi and guided with the usage. E.S. and F.M.J. provided Glioblastoma and HNSCC cell lines respectively and shared their p53 mutation status and HPV status. J-C.B. provided input about the p53 isoform data interpretations and manuscript writing. J.V.H. conceived the ideas and oversaw the research.

## Expanded View Figure legends

**Fig EV1. MDM2 copy number and mRNA expression status.**

(A) mRNA expression and MDM2 copy number analysis of NSCLC cell lines

(B) Western blot showing MDM2 expression in GBM cell lines shown.

(C) mRNA expression and MDM2 copy number analysis of HNSCC cell lines.

Expression analysis done using RNAseq data. HPV status and *TP53* mutation staus are indicated. CN, copy number. Arrows indicate the cell lines used in this study.

**Fig EV2. NMD inhibition prolongs p53β/γmRNA decay**

(A) mRNA decay analysis of p53α, p53β and p53γ mRNA transcripts from either DMSO or NMDi (1μM) treated A549 cells. NMDi and 5,6-Dichloro-1-β-D-ribofuranosylbenzimidazole (100μM) treatment was done as described in Methods. RT-qPCR analysis shown are mean ± s.e of three independent experiments.

**Fig EV3. p53γ contributes to the increased p53 pathway activation upon NMD inhibition in HPV^+^ HNSCC cell line.**

(A) p53β and p53γ mRNA expression FC upon NMDi treatment in HMS001 cells. Cells were treated with the indicated siRNAs for 54 hours and then with either DMSO (control) or 1μM NMDi, for 16 hours. RT-qPCR (n = 2 technical repeats) shown are representatives of two independent experiments. Mean ± s.e., p values, two tailed t-tests, *≤0.05, **< 0.01.

(B) Western analysis showing p53α (full-length) and truncated p53β/γ expression. siRNA and NMDi treatment were done as in (A). Western analysis shown are representatives of two independent experiments.

(C, D and E) mRNA expression FC of p53 transcriptional targets p21 (C), GADD45A (D) and PUMA (E). siRNA and NMDi treatment were done as in (A). RT-qPCR (n = 2 technical repeats) shown are representatives of 3 independent experiments. Mean ± s.e., p values, two tailed t-tests, *≤0.05, **< 0.01, ***< 0.001.

(Fand G) Western analysis showing p21 (F) and PUMA (G) expression. siRNA and NMDi treatment were done as in (A). Lower panels in F and G are the quantifications (2 technical repeats) of the Western blots, normalized to Vinculin. Western analysis shown are representatives of two independent experiments. Mean ± s.e., p values, two tailed t-tests, *≤0.05, **< 0.01, ***< 0.001.

**Fig EV4. p53 function restoration strategy by NMD inhibition in transcripts bearing mutations downstream of exon 9.**

(A) In the absence of NMD inhibition, NMD-inducing nonsense or splice site mutations downstream of p53 exon 9 cause mRNA degradation

(B) By inhibiting NMD, p53 transcripts bearing NMD-inducible mutations downstream of exon 9 can be protected and translated into near full-length functional proteins.

(C) Missense mutations downstream of exon 9 can generate mutant p53 protein with LOF/GOF alteration.

(D) NMD inhibition can potentially overcome the effect of LOF/GOF mutations downstream of exon 9 by enabling the expression of functional p53β/γ which are C-terminally truncated.

**Fig EV5. NMDi reduces tumor cell viability, disrupts cell cycle progression, reduces tumor growth and increases p53γ expression in tumors.**

(A) NSCLC, HNSCC and GBM cell lines are sensitive to NMDi treatment. NMDi treatment and IC50 calculations were done as described in Methods.

(B) HPV^+^ HNSCC cell lines are more sensitive to NMDi than indicated chemotherapy agents and nutlin. Cell lines shown were treated with the agents shown. IC50 shown are average from three technical replicates. Experiment was repeated two separate times.

(C) NMDi disrupts cell cycle progression. A549 cells were treated with either DMSO or NMDi (1μM) for the indicated time points, fixed in 70% ethanol, stained with propidium iodide and analyzed by Flow Cytometry as described in Methods.

(D) Orthotopic tumor growth model of UMSCC47 on nude mice tongue. Mice were injecred with UMSCC47 cells and randomized as described in the Methods. Randomized mice (n=10 per group) were treated either with vehicle or with 50mg/kg of NMDi for indicated number of days on a five days of treatment and two days of no treatment regimen.

(E) mRNA expression FC in vehicle or NMDi treated tumor tissue samples derived from A549 or UMSCC47 subcutaneous xenografts. Tumors were harvested at the end of the treatment, mRNA expression assessed by RT-qPCR. Mean ± s.e., n = 5 each.

